# Preconditioning by voluntary wheel running attenuates later neuropathic pain via Nrf2 antioxidant signaling in rats

**DOI:** 10.1101/2021.07.21.452532

**Authors:** Suzanne M. Green-Fulgham, Michael E. Harland, Jayson B. Ball, Heather D’Angelo, Renee A. Dreher, Jiahe Li, Michael J. Lacagnina, Sabina A. Lorca, Andrew J. Kwilasz, Steven F. Maier, Linda R. Watkins, Peter M. Grace

## Abstract

Animal and human studies have shown that exercise prior to nerve injury prevents later chronic pain, but the mechanisms of such preconditioning remain elusive. Given that exercise acutely increases formation of free radicals, triggering antioxidant compensation, we hypothesized that voluntary running preconditioning would attenuate neuropathic pain by supporting redox homeostasis after sciatic nerve injury in male and female rats. We show that 6 weeks of voluntary wheel running suppresses neuropathic pain development induced by chronic constriction injury (CCI) across both sexes. This protection was associated with reduced nitrotyrosine immunoreactivity—a marker for peroxynitrite—at the sciatic nerve injury site. Our data suggest that prior voluntary wheel running does not reduce production of peroxynitrite precursors, as expression levels of inducible nitric oxide synthase and NADPH oxidase 2 were unchanged. Instead, voluntary wheel running increased superoxide scavenging by elevating expression of superoxide dismutases 1 and 2. Prevention of neuropathic pain was further associated with activation of the master transcriptional regulator of the antioxidant response, nuclear factor E2-related factor 2 (Nrf2). Six weeks of prior voluntary wheel running increased Nrf2 nuclear translocation at the sciatic nerve injury site; in contrast, 3 weeks of prior wheel running, which failed to prevent neuropathic pain, had no effect on Nrf2 nuclear translocation. The protective effects of prior voluntary wheel running were mediated by Nrf2, as suppression was abolished across both sexes when Nrf2 activation was blocked during the running phase. This study provides insight into the mechanisms by which physical activity may prevent neuropathic pain.

## Introduction

Exercise can relieve established chronic pain in a range of preclinical models and clinical conditions (Dobson et al., 2014; Sterling et al., 2018; Guo et al., 2019). For example, preclinical and clinical studies have shown that exercise alleviates neuropathic pain (Cobianchi et al., 2010; Stagg et al., 2011; Li and Hondzinski, 2012; Groover et al., 2013; Benson et al., 2015; Bobinski et al., 2015; Grace et al., 2016b; Chhaya et al., 2019), a chronic pain condition caused by lesion or disease of the somatosensory system with limited pharmacotherapeutic options (Finnerup et al., 2015). In contrast to prior reports, we and others have shown that voluntary wheel running that ends prior to nerve injury protects against subsequent neuropathic pain in rodents (Grace et al., 2016b; Slivicki et al., 2019). Strikingly, protection was maintained for months after injury despite rats having no further running wheel access (Grace et al., 2016b). Supported by clinical literature, these findings have important public health implications as they suggest that an active lifestyle may prevent neuropathic pain (Landmark et al., 2011; Landmark et al., 2013). However, mechanisms by which exercise preconditioning prevents later neuropathic pain are unknown.

One hypothesis is that preconditioning prevents disbalance of redox homeostasis following peripheral nerve injury. That is, prior voluntary running may attenuate endogenous production of reactive oxygen and nitrogen species (ROS/RNS) after injury. For example, nuclear factor κB (NFκB) and mitogen activated protein kinases (MAPKs) upregulate NADPH oxidases (NOX) and nitric oxide synthases (NOS) (Anrather et al., 2006; Guo et al., 2007; Yoo et al., 2008). Attenuated NFκB and MAPK activation by prior voluntary running (Grace et al., 2016b) may therefore reduce NOX and NOS expression after peripheral nerve injury. Alternatively, prior voluntary running may prevent disbalance of redox homeostasis by increasing scavenging of ROS/RNS after injury. Repeated bouts of exercise acutely increase ROS production by skeletal muscle, leukocytes, and other cells (Margaritelis et al., 2020). In response, exercise increases antioxidant gene expression and protein levels (e.g., superoxide dismutase, glutathione) in numerous tissues (Done and Traustadottir, 2016). Attenuated ROS/RNS production, increased scavenging of ROS/RNS, or a combination of both are anti-nociceptive, given the causal role of unchecked ROS/RNS in neuropathic pain (Twining et al., 2004; Bonnefous et al., 2009; Kim et al., 2010; Doyle et al., 2012; Kallenborn-Gerhardt et al., 2012; Little et al., 2012; Hassler et al., 2014; Kallenborn-Gerhardt et al., 2014; Grace et al., 2016a; Shim et al., 2019). ROS/RNS and their reactive products promote neuronal hyperexcitability through a range of mechanisms, including increased activity of ion channels expressed by sensory neurons (e.g., transient receptor potential A1 and V1) (Trevisani et al., 2007; Patwardhan et al., 2009; Taylor-Clark et al., 2009; Shepherd et al., 2018), disrupted spinal GABAergic signaling (Yowtak et al., 2011; Yowtak et al., 2013), mitochondrial dysfunction (Schwartz et al., 2009; Janes et al., 2013; Li et al., 2020), and cytokine induction (Matata and Galinanes, 2002; Kamata et al., 2005; Doyle et al., 2012; Li et al., 2020). Based on these lines of evidence, we hypothesized that voluntary running preconditioning would suppress the development of neuropathic pain by supporting redox homeostasis after peripheral nerve injury.

Our study shows that voluntary running preconditioning increases antioxidant compensation at the nerve injury site, independently of sex. We also provide evidence that prevention of neuropathic pain is mediated by exercise-induced activation of nuclear factor E2-related factor 2 (Nrf2)—the master transcriptional regulator of the antioxidant response (Cuadrado et al., 2019; Grace et al., 2021). These findings offer insight into the mechanisms by which prior voluntary exercise controls redox balance after injury, guarding against the development of chronic pain in both males and females.

## Materials and Methods

### Animals

Pathogen-free adult male and female Sprague Dawley rats (Envigo, Indianapolis, IN), 10 weeks old on arrival were used in all experiments. Rats were same sex pair housed on arrival in temperature-controlled (23 ± 3°C) and light-controlled (12 hours light–dark cycle; lights on at 07:00 hours) rooms with standard rodent chow and water available *ad libitum*. Housing conditions during wheel running are described below. All procedures were approved by the University of Colorado Boulder and MD Anderson Cancer Center Animal Care and Use Committees.

### Voluntary wheel running

Rats were single-housed with in-cage running wheels, whereas sedentary control animals were single-housed in standard cages. Single housing allowed for accurate running data collection and avoided competition for the wheel. We have previously shown that there are no differences in sensory thresholds between sedentary rats housed with a locked wheel versus a standard homecage, either prior to or across time after nerve injury (Grace et al., 2014a; Grace et al., 2016b; Green-Fulgham et al., 2020). Rats were given unrestricted access to in-cage running wheels, continuously for the 3 or 6 weeks immediately before surgery. Voluntary wheel running concluded on the day of surgery, when rats were returned to the home cage and standard pair housing with the previous cage mate was reinstated for all subjects. Daily wheel revolutions and running speed (meters/min) were recorded digitally using Scurry activity monitoring software by Lafayette Instruments (Lafayette, IN), or by Vital View software (Bend, OR), and the weekly distance traveled was calculated by multiplying the number of revolutions by the wheel circumference (1.081 m).

### Chronic constriction injury (CCI) surgery

Neuropathic pain was induced using the sciatic nerve CCI model (Bennett and Xie, 1988). Briefly, surgery was performed under isoflurane anesthesia at the mid-thigh level of the left hind leg. The shaved skin was treated with Nolvasan, and the surgery was performed aseptically. The sciatic nerve was gently isolated, and four chromic gut sutures (cuticular 4-0 WebGut; Patterson Veterinary, Devens, MA) were loosely tied around the nerve. For sham surgery, the sciatic nerve was isolated, but no chromic gut sutures were tied around the nerve. Animals were monitored postoperatively until fully ambulatory before returning them to their home cage.

### Mechanical allodynia

Testing was conducted blind with respect to group assignment, and results were confirmed by at least 2 different investigators in each experiment. Rats received at least three 60-minute habituations to the test environment before behavioral testing. The von Frey test (Chaplan et al., 1994) was performed as previously described in detail (Chacur et al., 2001; Milligan et al., 2001) within the sciatic innervation region of the hind paws. Assessments were made before voluntary wheel running, before CCI surgery, and then at regular intervals. A logarithmic series of 10 calibrated Semmes–Weinstein monofilaments (von Frey hairs; Stoelting, Wood Dale, IL) were applied to the hind paws to define the threshold stimulus intensity required to elicit a paw withdrawal response. Log stiffness of the hairs ranged from manufacturer-designated 3.61 (0.40 g) to 5.18 (15.14 g) filaments. The behavioral responses were used to calculate the absolute threshold (the 50% probability of response) by fitting a Gaussian integral psychometric function using a maximum-likelihood fitting method (Harvey, 1986; Treutwein and Strasburger, 1999) as described previously (Milligan et al., 2000; Milligan et al., 2001). This fitting method allowed parametric analyses that otherwise would not be statistically appropriate (Harvey, 1986; Treutwein and Strasburger, 1999).

### Pharmacological Nrf2 inhibition

Male and female rats were administered trigonelline (Cayman Chemical, Ann Arbor, MI) in the drinking water to inhibit Nrf2 activation for the 6 week duration of wheel running (Boettler et al., 2011; Li et al., 2020). No trigonelline was administered at any time post CCI surgery. Trigonelline concentrations were based on the average water consumption for each rat from the prior week, to achieve average doses of 100 mg/kg (made fresh daily). We selected a pharmacological approach as it allows temporal control over inhibition of Nrf2, specifically during the period of voluntary wheel running. We did not include sham controls, as we have previously shown that trigonelline does not alter sensory thresholds in sham-operated animals (Li et al., 2020). Regular drinking water was used as vehicle control.

### Tissue collection

On days 7 and 14 post CCI/sham surgery, rats were overdosed with sodium pentobarbital, and then transcardially perfused with ice-cold saline. Ipsilateral sciatic nerves, medial to the injury site, L4/5 dorsal root ganglia (DRG), and L4/5 spinal cords were isolated. For PCR and Western blot, tissues were snap frozen in liquid nitrogen and stored at −80 °C until use. For immunohistochemistry, the saline perfusion was followed by ice-cold 4% paraformaldehyde in 0.1 M phosphate buffer (pH 7.4).

### Immunohistochemistry

Tissues were postfixed by immersion in 4% paraformaldehyde for 24 hours4 °C, and cryoprotected sequentially in 15%, 20%, and 30% sucrose with 0.1% azide at 4°C. Sections were then freeze-mounted in OCT, and frozen sections were cut at 12 μm (sciatic nerves), 10 μm (DRG), or 20 μm (spinal cord). For analysis of nitrotyrosine, sections were washed, permeabilized with 0.3% hydrogen peroxide, blocked for 1 hour with 10% NGS, 0.3% Triton-X in phosphate buffered saline (PBS), and then incubated overnight at 4°C in 2% normal goat serum together with mouse anti-nitrotyrosine (1:150; Cayman Chemical). Slides were then washed and incubated in goat anti-mouse biotin secondary antibody (1:200; Jackson Immuno Research, West Grove, PA) for 2 hours. Sections were washed, incubated in ABC solution (Vector Laboratories, Burlingame, CA) for 2 hours, washed again, and incubated in inactive DAB (Sigma) for 10 minutes. DAB was then activated with beta-D glucose (10 mg/mL) and slides were incubated for 10 minutes, washed, and dried overnight. Slides were then dehydrated in increasing concentrations of ethanol (50%, 70%, 95%, and 100%), cleared in Citrisolv, dried and covered with DPX mountant (Sigma). Images were acquired using an Olympus BX61 microscope (Olympus, Center Valley, PA) with Cell Sens Dimension software (Olympus).

Images were converted to 32 bits and corrected for threshold, and densitometry analysis was performed using NIH Fiji software by an investigator who was blinded to treatment groups. For lumbar dorsal horn, images from four sections per animal were acquired at 20x magnification and the ipsilateral dorsal horn was selected for analysis. For sciatic nerve and DRG, images from four sections per animal were acquired in mosaic format with a mechanical stage at 40x magnification. Data are expressed as total area positive for staining within the region of interest.

For analysis of nuclear translocation of Nrf2, antigen retrieval was performed by incubation in citrate buffer at 95ºC for 15 minutes. Sections were then washed, blocked for 1 hour with 10% NGS, 0.3% Triton-X in PBS, and then incubated overnight at 4°C for 24 hours in 2% normal goat serum together with rabbit anti-Nrf2 at 1:500 (Abcam). Slides were then washed and incubated in the secondary antibody (1:1500; goat anti-rabbit, ThermoFisher Scientific, Waltham, MA) for 2 hours at room temperature, followed by 5 minute incubation in DAPI (1:5,000; ThermoFisher Scientific, Waltham, MA). Slides were then washed, dried, and covered with Prolong Gold mountant (Thermo Fischer Scientific; Waltham, MA). For each of sciatic nerve, DRG, and spinal cord, images from four sections per animal were acquired at 40x on a Nikon Eclipse Ti laser scanning confocal microscope, using Nikon Elements software and converted to maximum intensity composite images using NIH Fiji software. Nuclear translocation of Nrf2 was determined by calculating the percentage of DAPI^+^ nuclei that were colocalized with Nrf2, relative to total number of DAPI^+^ nuclei.

### Real-time polymerase chain reaction

RNA was isolated from frozen tissues using the phenol: chloroform extraction method. RNA concentrations > 100 ng and purity > 1.6 A260/A280 for cDNA normalization were confirmed using a nanodrop spectrophotometer (Thermo Fischer Scientific; Waltham, MA). The samples were stored at −80°C until use. cDNA synthesis was performed using Superscript IV (Invitrogen; Carlsbad, CA). RT-PCR was conducted using QuantiTect SYBR Green PCR Kit (Qiagen; Germantown, MD) in 96 well PCR plates (Bio-Rad) on a CFX96 Touch Real Time PCR Detection System (Bio-Rad). Primers were designed using GenBank (National Center for Biotechnology Information; http://www.ncbi.nlm.nih.gov), with sequences as follows: *Bactin* (forward: TTCCTTCCTGGGTATGGAAT; reverse: GAGGAGCAATGATCTTGATC); *Sod1* (forward: TGAAGAGAGGCATGTTGGAG; reverse: TCATCTTGTTTCTCGTGGAC); *Sod2* (forward: GAACCCAAAGGAGAGTTGCT; reverse: CTCCTTATTGAAGCCAAGCC); *Cybb* (forward: GGATGAATCTCAGGCCAATC; reverse: GGTGTTGACTTGCAATGGTC); *Nox4* (forward: TTCTCAGGTGTGCATGTAGC; reverse: CGGAACAGTTGTGAAGAGAAGC); *Nos2* (forward: ACCCAAGGTCTACGTTCAAG; reverse: GCTTCTTCAAAGTGGTAGCC); *HO1* (forward: CTATCGTGCTCGCATGAAC; reverse: CAGCTCCTCAAACAGCTCAA); *HO2* (forward: CCATACCAAAATGGCAGACC; reverse: CAAGGGCTGAGTATGTGAAG). Threshold for detection of PCR product was set in the log-linear phase of amplification and the threshold cycle (CT) was determined for each reaction. Each sample was measured in duplicate and the results were normalized to the expression of the housekeeping gene (β-actin), using the ΔΔCT method. β-actin expression was not different between treatment groups.

### Western blotting

Nuclear fractions from ipsilateral sciatic nerve injury sites from each rat were isolated with a NE-PER Nuclear and Cytoplasmic Extraction Kit (ThermoFisher Scientific), according to manufacturer’s instructions. Western blotting was performed as previously described (Grace et al., 2016b; Grace et al., 2016c; Li et al., 2020). In brief, extracted nuclear proteins were subjected to NuPAGE Bis-Tris (4 to 12%) gel electrophoresis under reducing conditions (ThermoFisher Scientific) and then electrophoretically transferred to nitrocellulose membranes (Bio-Rad, USA). Nonspecific binding sites on the membrane were blocked with Superblock buffer containing 0.1% Tween-20, 0.05% Tris-Chloride, and 0.03% 5 M NaCl for 1 h at 22° to 24°C. Membranes were subsequently incubated overnight at 4°C with primary anti-Nrf2 antibody (1:1,000; rabbit polyclonal IgG; Abcam) and antihistone H3 antibody (1:2,000; rabbit polyclonal IgG; Abcam) (loading control) in blocking buffer containing 0.1% Tween-20. The membranes were then washed with PBS containing 0.1% Tween-20, and probed with horseradish peroxidase secondary antibody (1:5,000; goat polyclonal IgG; Jackson ImmunoResearch, USA) in blocking buffer containing 0.1% Tween-20 for 1 h at 22° to 24°C. After washing with PBS containing 0.1% Tween-20, membranes were developed with enhanced chemiluminescent substrate (ThermoFisher Scientific), and scanned on an ImageQuant LAS 4,000 mini (GE, USA). Densitometry analysis was performed using ImageQuant TL software (GE). Data were normalized to loading control (histone H3).

### Statistics

Mechanical allodynia was analyzed as the interpolated 50% thresholds (absolute threshold). Where appropriate, one-way ANOVAs or unpaired t tests were used to confirm that there were no baseline differences in absolute thresholds between treatment groups. Differences in mechanical allodynia between treatment groups following CCI were determined using two-way repeated measures ANOVA followed by Sidak’s post hoc tests where appropriate. Differences in male and female running behavior was determined by two-way repeated measures ANOVA followed by Sidak’s post hoc tests where appropriate. Differences in protein and gene expression were determined by three-way ANOVA, with sex, wheel running and injury as variables, followed by Tukey’s post hoc tests where appropriate. If no there were no main effects of sex, then data from males and females within the same treatment groups were consolidated. Correlations were determined by Pearson’s correlation, with a Bonferroni correction for multiple comparisons. All analyses were performed using Prism 9 (GraphPad). Results are expressed as mean ± SD. P<0.05 was considered statistically significant.

## Results

### Voluntary running preconditioning attenuates subsequent neuropathic pain in female rats

We have previously shown that six weeks of voluntary wheel running prior to CCI attenuates subsequent mechanical allodynia in male rats (Grace et al., 2016b), but it is not yet known if such preconditioning similarly prevents neuropathic pain in female rats. To address this question, female rats were housed in identical running conditions as males in prior experiments, with unlimited access to in-cage running wheels for six weeks prior to CCI, after which they were returned to their home cage. Prior voluntary wheel running attenuated mechanical allodynia after CCI in female rats (Fig. 1A; treatment x time: F_51,544_ =8.290, p<0.0001; treatment F_3,32_= 185.6, p<0.0001; time F_17,544_ = 32.76, p<0.0001), from weeks 2-7 (p<0.05) and 9-13 after injury (p<0.05). Protection from neuropathic pain in female rats was similar in magnitude to what we previously reported in male rats (reproduced in Fig. 1B) (Grace et al., 2016b). Thus, six weeks of voluntary wheel running prior to CCI protects against subsequent neuropathic pain in both sexes.

**Figure 1.**
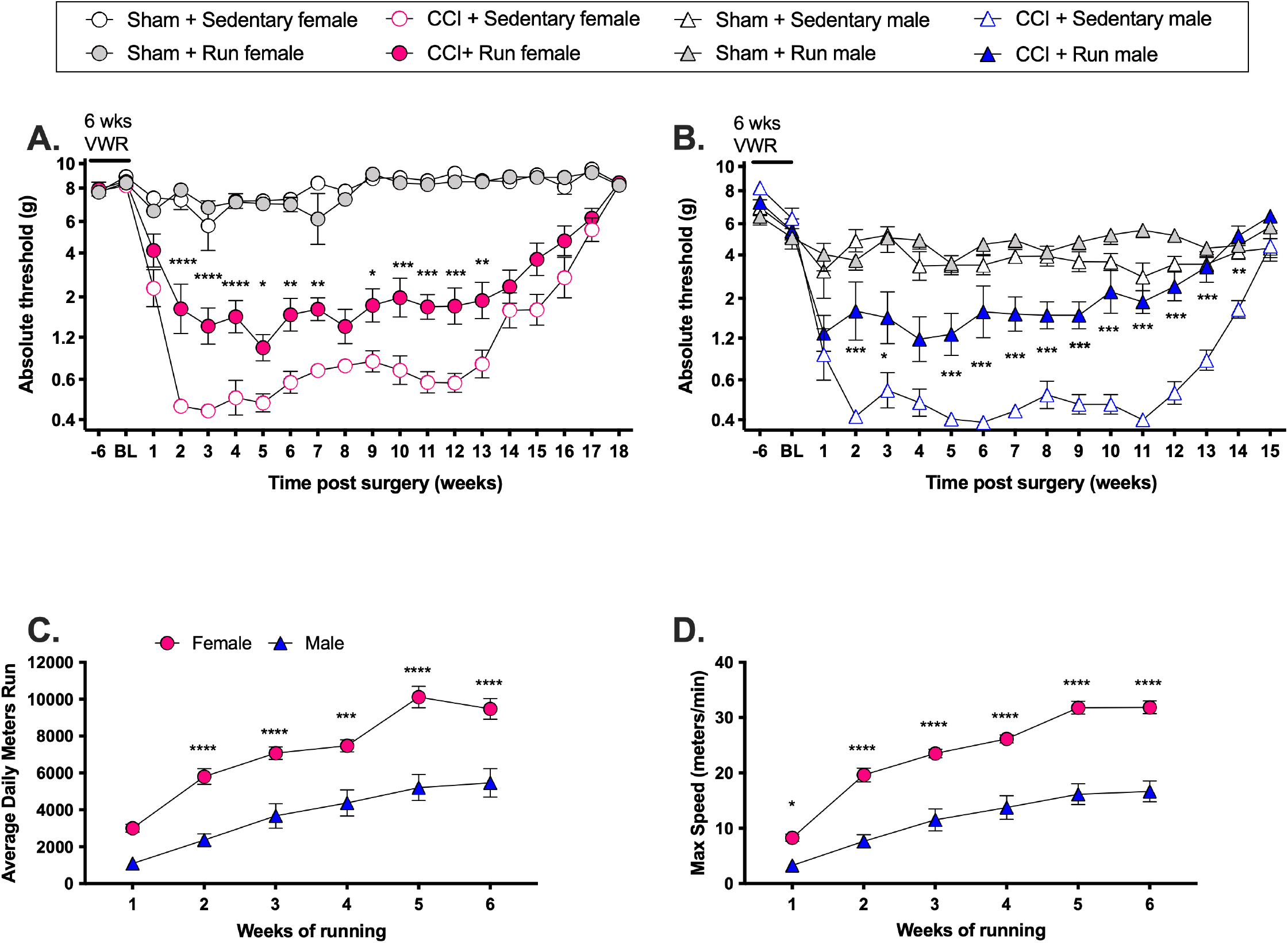
**(A)** Six weeks of prior voluntary wheel running (VWR) attenuates subsequent chronic constriction injury (CCI)-induced allodynia in female rats, and **(B)** male rats (historic data, (Grace et al., 2016b)). Rats were single-housed with an in-cage running wheel (run) for 6 weeks, or in standard housing (sedentary) for 6 weeks before sham or CCI surgeries were performed. Rats were returned to standard paired housing with previous cage mates after surgery. Von Frey thresholds were assessed before and after 6 weeks of exercise, and at weekly intervals after CCI. *p<0.05, **p<0.01, ***p<.001, ****p<.0001 CCI + sedentary vs. CCI + run. Female rats ran greater distances **(C)** with greater maximum speeds **(D)** than male rats. Rats were single-housed with an in-cage running wheel for 6 weeks. Running distance and speed were continuously recorded. Note y axis differs between graphs. *p<0.05, ***p<.001, ****p<.0001, male vs. female.

### Influence of voluntary wheel running parametrics on prevention of neuropathic pain

There were considerable differences in running activity between individual rats and across sexes (Fig 1C, D) as we have reported previously (Grace et al., 2016b). Weekly distances traveled in the final week of running ranged from 5,349 to 12,639 m in females, and 1,037 to 9,991 m in males. Maximum running speeds ranged from 23 to 40 m/min in females, and 3 to 26 m/min in males. However, these differences in running parametrics had little bearing on the attenuation of allodynia, as sensory thresholds did not correlate with distances travelled (females: p=0.8395, r= −0.0792; males: p=0.9087, r= 0.0610, week 2 post CCI) or maximum running speeds (females: p=0.1291, r= −0.5450, week 2 post CCI, (data not collected for males)) at any time point post CCI. Although female rats ran greater distances (Fig. 1C) and at greater speeds (Fig. 1D) than male rats, congruent with previous reports (Eikelboom and Mills, 1988), these differences did not influence sensory thresholds; the magnitude by which prior voluntary wheel running attenuated allodynia was statistically indistinguishable between the sexes (male vs female F_1,13_= 0.02, p=0.8936; Figs 1A, B). Furthermore, running speeds or distances did not correlate with any biochemical measures presented below.

We next asked whether a shorter duration of voluntary wheel running prior to CCI would still confer protection against subsequent neuropathic pain. Three weeks of voluntary wheel running was not sufficient to protect rats of either sex from neuropathic pain (Fig. 2; treatment x time: F_3,66_ =0.74, p=0.5279; treatment: F_1,22_= 0.62, p=0.4398; time: F_2.47,54.3_= 46.70, p<0.0001). Together, these results point to a low threshold of weekly distances travelled for both male and female rats, beyond which, neuropathic pain is not additionally attenuated. Notably, the robust benefits of preconditioning for prevention of neuropathic pain start between three and six weeks of voluntary wheel running.

**Figure 2.**
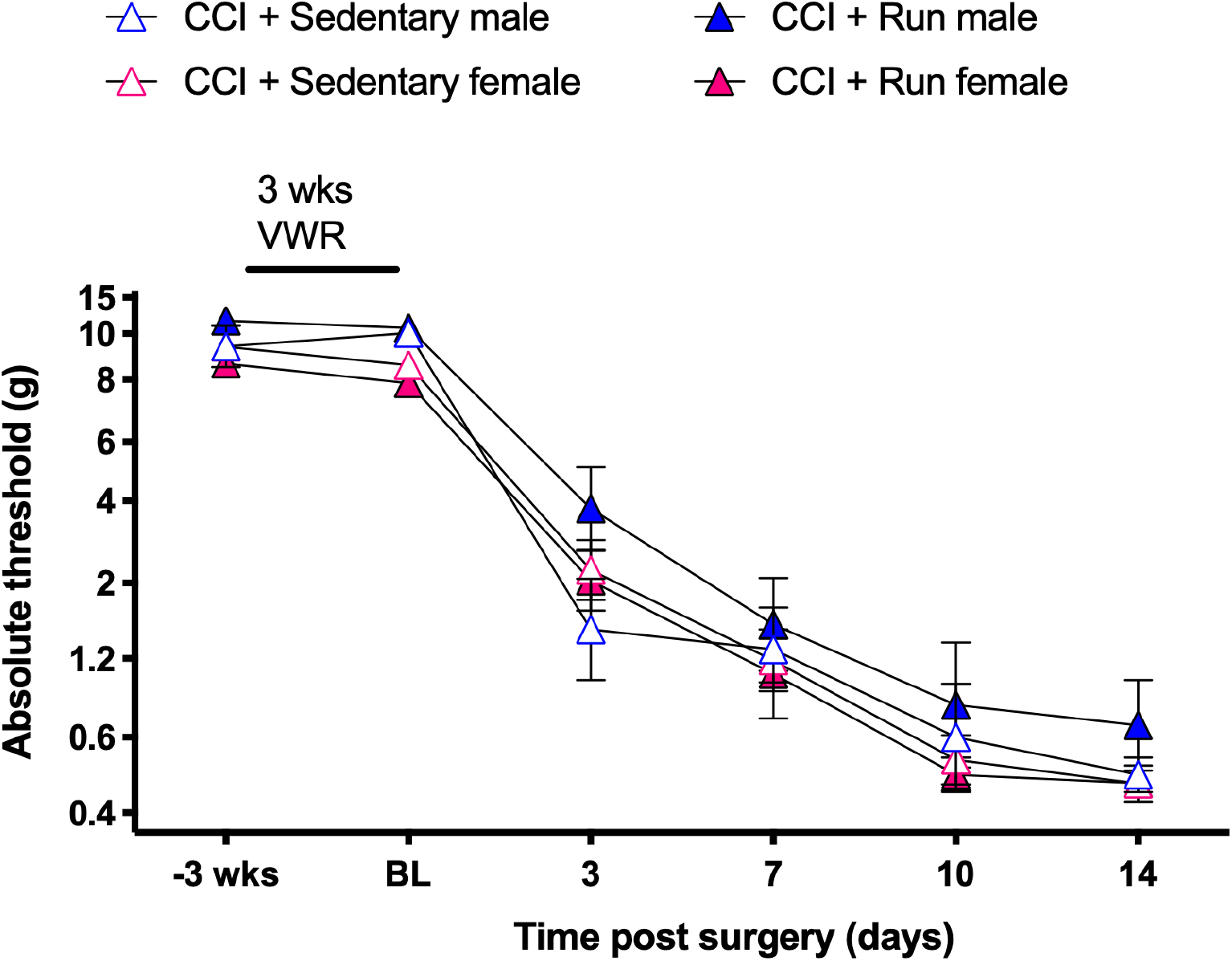
Three weeks of prior voluntary wheel running (VWR) does not attenuate subsequent chronic constriction injury (CCI)-induced allodynia in female and male rats. Rats were single-housed with an in-cage running wheel (run) for 3 weeks, or single-housed in standard housing (sedentary) for 3 weeks before CCI surgeries were performed. Rats were returned to standard housing with previous cage mates after surgery. Von Frey thresholds were assessed before and after 3 weeks of exercise, and at 3-4 day intervals after CCI.

### Prior voluntary running reduces nitrotyrosine levels at the site of sciatic nerve injury

It is well appreciated that peripheral nerve injury disrupts redox homeostasis in the pain neuraxis, which is causal to neuropathic pain (Bonnefous et al., 2009; Kim et al., 2010; Salvemini et al., 2011; Doyle et al., 2012; Kallenborn-Gerhardt et al., 2012; Little et al., 2012; Grace et al., 2014b; Kallenborn-Gerhardt et al., 2014; Grace et al., 2016a; Shepherd et al., 2018; Shim et al., 2019). To test our hypothesis that prior exercise would restore redox homeostasis after peripheral nerve injury, we first investigated peroxynitrite—a reactive product of nitric oxide and superoxide, and an exemplar reactive species that is causal to neuropathic pain (Salvemini et al., 2011; Doyle et al., 2012; Little et al., 2012; Janes et al., 2013). Tyrosine nitration (nitrotyrosine) was quantified as an indirect measure of peroxynitrite (Wang et al., 2004). Nitrotyrosine immunoreactivity was increased at the sciatic nerve injury site at days 7 and 14 post CCI, compared to sham surgery (day 7 males and females p<0.0001; day 14 sedentary males and females p<0.0001) (Fig. 3A,B). Both sexes exhibited increases, though nitrotyrosine elevation was greater in females compared to males at day 14 post CCI (p=0.0076). Voluntary wheel running prevented these increases in both sexes at day 7 (males and females p<0.0001) and day 14 (females: p<0.0001; males: p=0.0293). There were no correlations between running wheel activity (distance travelled of speeds) and nitrotyrosine levels in the sciatic nerve (female day 7: p=0.3514, r=-0.4662; male day 7: p=0.1621, r=0.6502; female day 14: p=0.4599, r=-0.2362; male day 14; p=0.0223, r=0.8756). CCI also induced a significant increase in nitrotyrosine in the DRG at day 7 and day 14 in sedentary rats of both sexes (day 7: p<0.0001; day 14 p=0.0017), and in runners of both sexes at day 14 (p=0.011)(Fig. 3C,D), but voluntary running did not modify nitrotyrosine expression in DRG (interaction day 7: p=0.0779, day 14 p=0.9099) (Fig. 3C,D). Neither CCI nor voluntary running induced significant changes in nitrotyrosine immunoreactivity in the lumbar spinal cord at either timepoint or sex (interaction day 7: p=0.7979; day 14 p=0.1545) (Fig. 3E,F). These data show that voluntary wheel running prevents increases in peroxynitrite, an exemplar free radical, at the sciatic nerve injury site.

**Figure 3.**
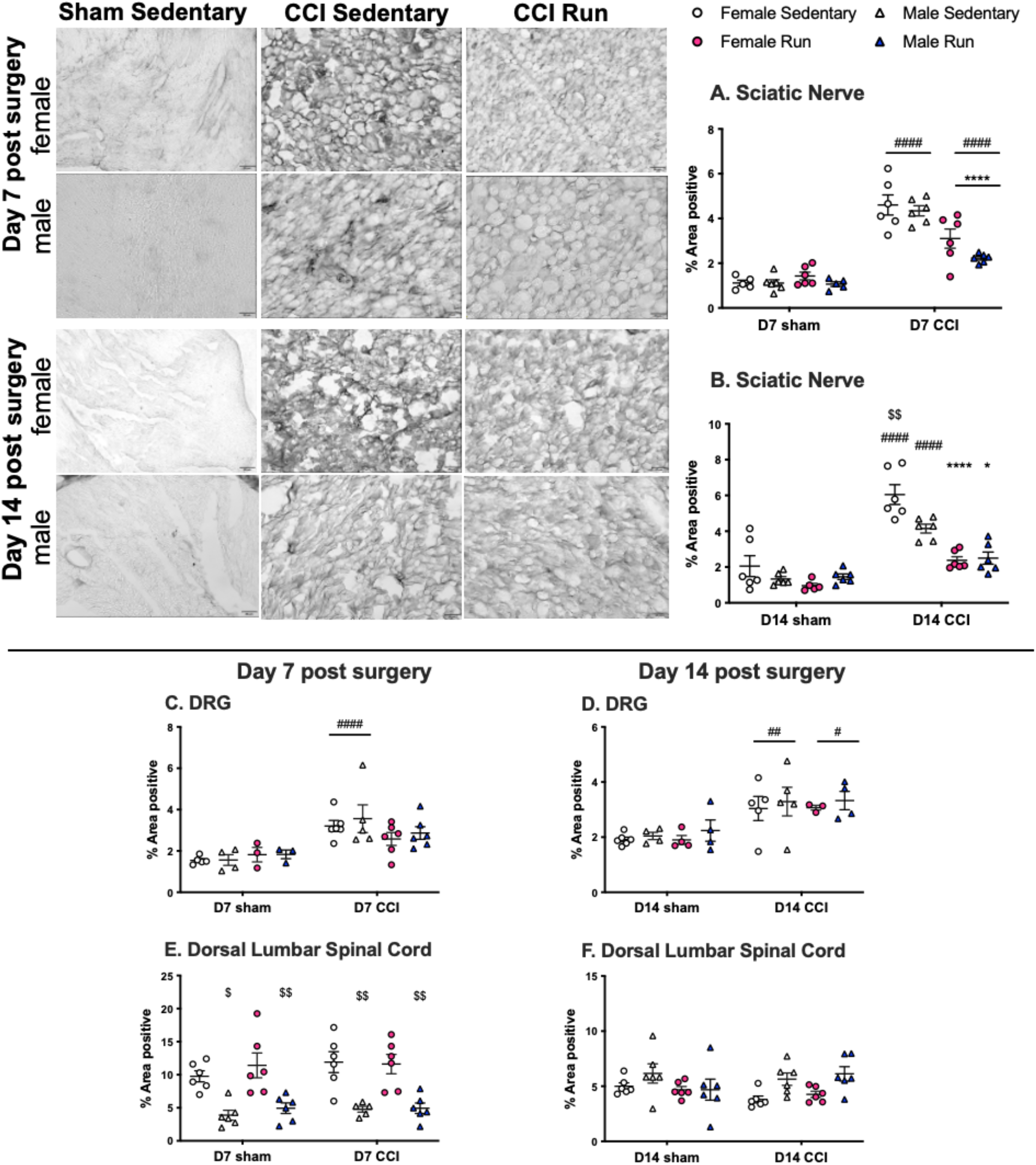
Nitrotyrosine protein levels were quantified at the medial sciatic injury site **(A,B)** scale bar = 20um, representative sedentary sham images shown as no difference was observed between sedentary shams and runner shams, ipsilateral L4/5 DRG **(C,D)**, and ipsilateral lumbar dorsal horn of the spinal cord **(E,F)** at day 7 and 14 post surgery. Note y axis differs between graphs. *p<0.05, ****p<.0001, CCI + sedentary vs. CCI + run. #p<0.05, ##p<0.01, ####p<.0001, sham vs. CCI. $p<0.05, $$p<0.01, male vs. female within each group. Males and females consolidated when no significant sex differences.

### Running preconditioning does not alter expression of iNOS and NADPH oxidases after nerve injury

There are at least two potential mechanisms by which prior voluntary running may reduce levels of peroxynitrite: attenuated nitric oxide/superoxide production, and/or increased antioxidant scavenging. To test these hypotheses, we first investigated expression of enzymatic generators of nitric oxide (iNOS) and superoxide (NOX isoforms). Since prior voluntary wheel running reduces activation of NFκB and MAPKs (Grace et al., 2016b), we reasoned that downstream induction of iNOS and NOX isoforms may also be reduced (Anrather et al., 2006; Guo et al., 2007; Li et al., 2007; Yoo et al., 2008). Expression of iNOS in the sciatic nerve was elevated to a similar extent in both sexes at days 7 and 14 after CCI (p<0.0001) (Fig 4 A,B). CCI also increased NOX2 expression in the sciatic nerve at day 7 (sedentary rats: p=0.0010; runners: p=0.0013) and at day 14 (sedentary and runner females: p<0.0001; male runners p=0.0215; sedentary males p=0.0512) (Fig 4 C,D). Although injury increased NOX2 expression in both sexes, females had a greater magnitude increase at day 14 (Fig 4D). Prior voluntary wheel running did not prevent increases in iNOS (day 7: p=0.9474; day 14: p=0.9985) or NOX2 (day 7: p=0.0728; day 14 females p=0.9998; day 14 males p=0.9838) expression at any timepoint or in either sex following CCI (Fig 4). There were no correlations between running distance or speeds and expression after CCI of iNOS (female day 7: p=0.8088, r=-0.1134; male day 7: p=0.513, r=0.3002; female day 14: p=0.6419, r=-0.2159; male day 14; p=0.1995, r=0.5072) or in NOX2 (female day 7: p=0.4899, r=-0.1934; male day 7: p=0.7258, r=-0.0951; female day 14: p=0.4003, r=0.2345; male day 14; p=0.8172, r=-0.0628).

**Figure 4.**
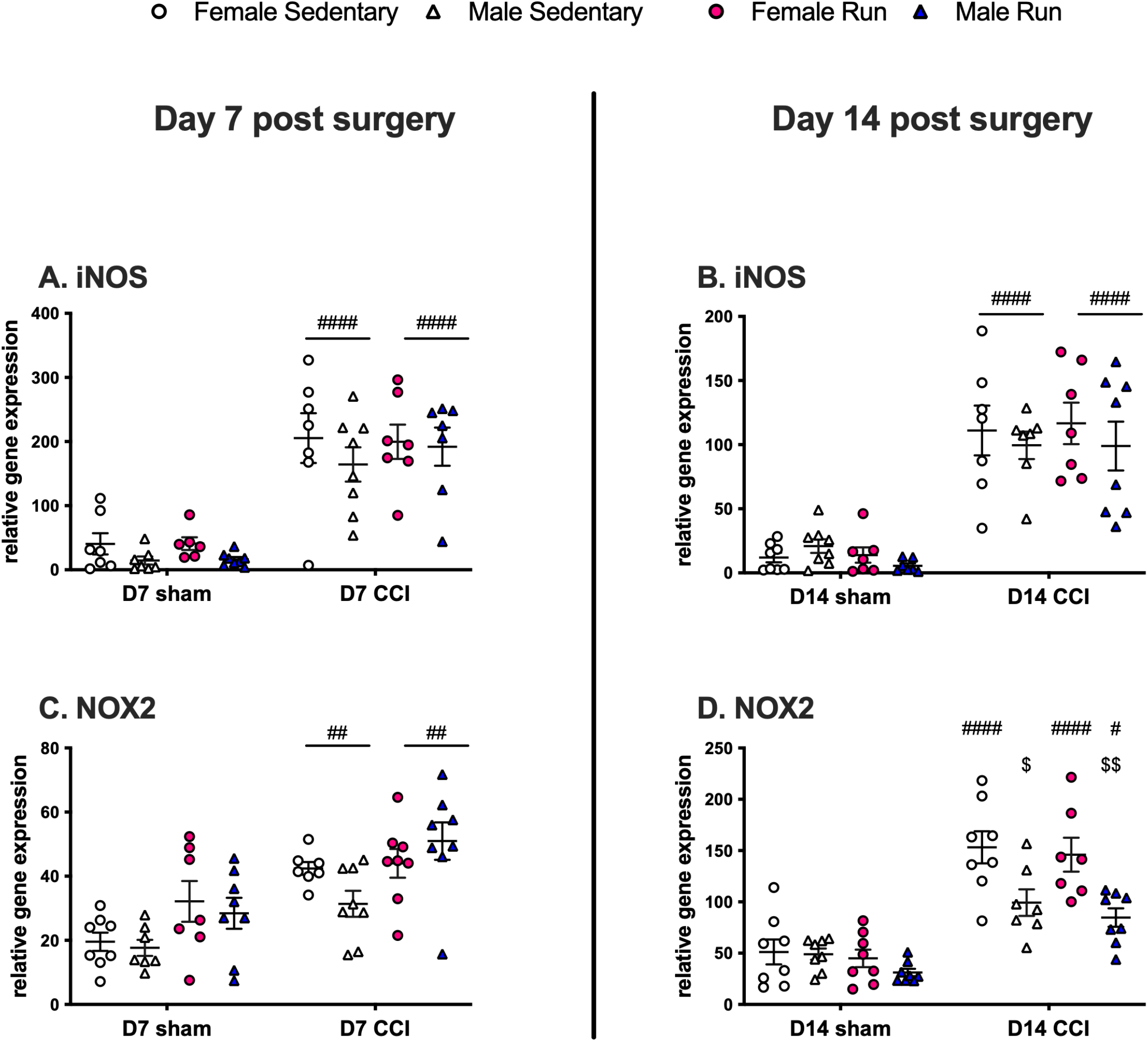
iNOS **(A,B)** and NOX2 **(C,D)** gene expression were quantified at the medial sciatic injury site at day 7 and 14 post surgery. Note y axis differs between graphs.**p<0.01, ****p<.0001. *p<0.05, **p<0.01, ***p<.001, ****p<.0001, CCI + sedentary vs. CCI + run. #p<0.05, ##p<0.01, ###p<.001, ####p<.0001, sham vs. CCI. $p<0.05, $$p<0.01, $$$p<.001, $$$$p<.0001, male vs. female within each group. Males and females consolidated when no significant sex differences.

Neither CCI nor voluntary running altered iNOS or NOX2 expression in the DRG at either timepoint, compared to sham and sedentary controls (iNOS day 7: p=0.8541, iNOS day 14: p=0.7782, NOX2 day 7: p=0.3916, NOX2 day 14: p=0.6446) (data not shown). In the spinal cord, neither CCI nor voluntary running altered iNOS expression at either timepoint, compared to sham and sedentary controls (day 7: p=0.2340, day 14 p=0.2539), but NOX2 was increased by CCI surgery at day 7 (sedentary males and females: p<0.0001, runners p=0.0017). Spinal NOX2 after CCI was decreased in rats with prior running at day 7 (p=0.0099) (data not shown). With some exceptions in constitutive expression levels, there were no differences in iNOS or NOX2 expression between the sexes in either tissue (data not shown). We also quantified NOX4 expression, but there were minimal effects of CCI and no changes due to running in any site or condition (sciatic: day 7: p=0.4374, day 14: p=0.6017; DRG: day 7: p=0.8764, day 14: p=0.3284; spinal cord: day 7: p=0.1537, day 14: p=0.4190)(data not shown). Based on these data, it is unlikely that running preconditioning attenuates peroxynitrite levels by reducing production of precursor nitric oxide and superoxide.

### Running preconditioning increases expression of superoxide dismutases after nerve injury

We next investigated increased antioxidant scavenging as the alternate path by which prior running could reduce levels of peroxynitrite. Superoxide dismutases (SOD1, SOD2) are the primary antioxidants that catabolize superoxide, while heme oxygenases (HO-1, HO-2) catalyze conversion of heme to nitric oxide scavengers and other antioxidants (Dore et al., 1999; Mancuso et al., 2006; Radi, 2018). Elevation of either or all antioxidants would eliminate critical substrates for peroxynitrite formation.

At day 7, prior running increased expression of cytosolic SOD1 and mitochondrial SOD2 in sciatic nerves compared to sedentary controls in both sexes (SOD1: sham operated: p<0.0001; CCI: p=0.0024; SOD2: sham operated: p=0.0035; CCI: p<0.0001) (Fig. 5A,C). CCI surgery decreased expression of SOD1 at day 7, whereas SOD2 expression was increased with CCI relative to sham (Fig. 5A,C) (SOD1: sedentary: p=0.0004, runners p<0.0001; SOD2: sedentary p=0.0022, runners p<0.0001).

**Figure 5.**
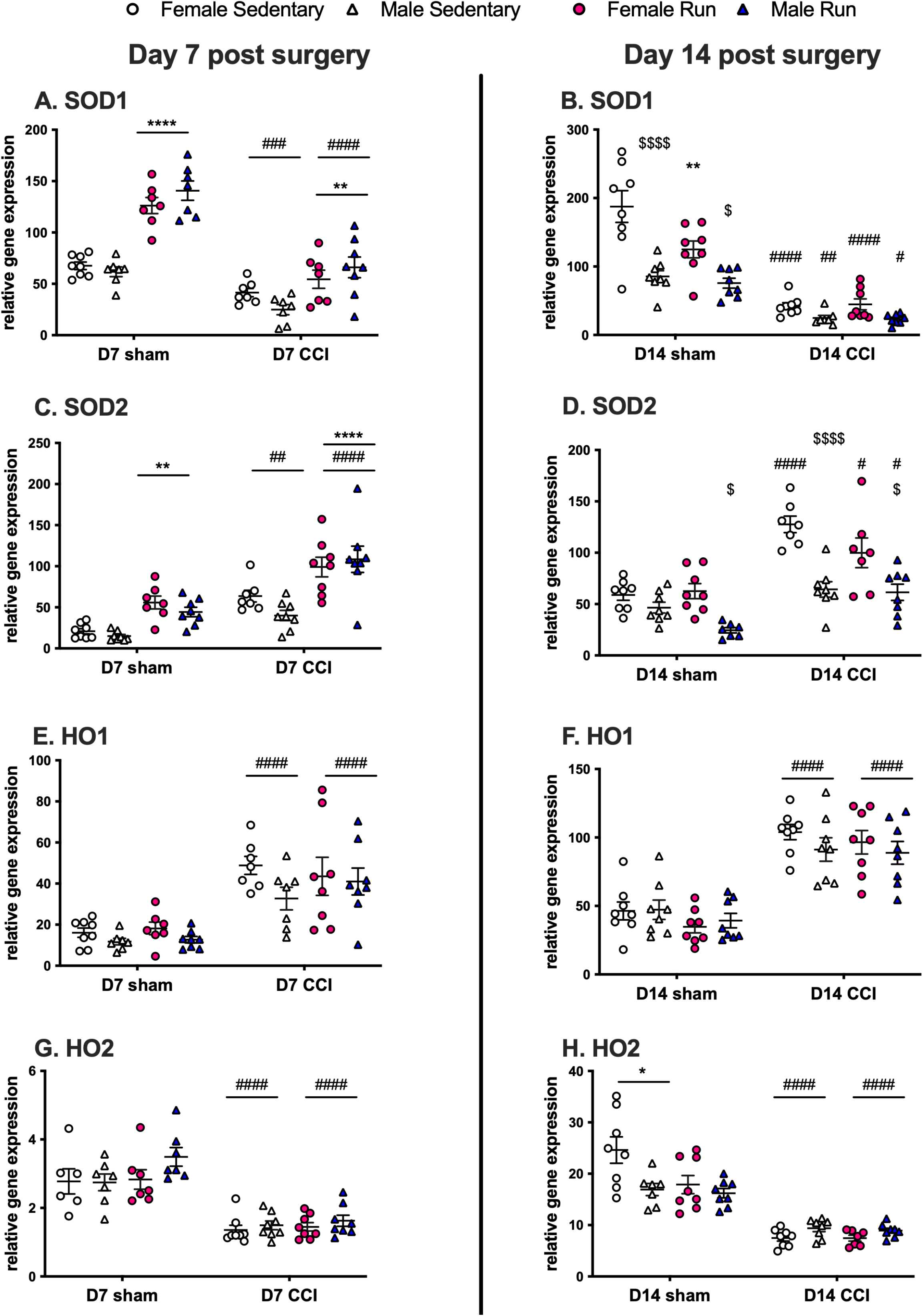
Superoxide dismutase 1 **(A,B)** and superoxide dismutase 2 **(C,D)** heme oxygenase 1 **(E,F)** and heme oxygenase 2 **(G,H)** gene expression were quantified in the medial sciatic injury site at day 7 and 14 post surgery. Note y axis differs between graphs. *p<0.05, **p<0.01, ****p<.0001, CCI + sedentary vs. CCI + run. #p<0.05, ##p<0.01, ###p<.001, ####p<.0001, sham vs. CCI. $p<0.05, $$$$p<.0001, male vs. female within each group. Males and females consolidated when no significant sex differences.

While SOD1 decreases and SOD2 increases were maintained after CCI at day 14 (SOD1: sedentary females: p<0.0001; female runners: p<0.0001; sedentary males: p=0.0073; male runners: p=0.0235; SOD2: sedentary females: p<0.0001; female runners: p=0.0291; male runners: p=0.0325), the effect of prior running on SOD2 expression was no longer apparent (Fig. 5B,D). SOD1 elevation in sham female runners was still apparent at day 14 (p=0.0033). Some sex differences in SOD expression were observed at day 14 post surgery, with expression remaining higher in female rats than males in all cases. SOD1 expression was higher in females than males in sham rats (p<0.0001 sedentary, p=0.0414 runners), and SOD2 expression was also higher in females (sedentary CCI p<0.0001; CCI runners p=0.0221; sham runners p=0.0248). There were no correlations between wheel running distance or speeds and sciatic expression levels after CCI of SOD1 (day 7 females: p=0.2364, r=0.5155; day 7 males: p=0.9969, r=-0.0016; day 14 females: p=0.4287, r=0.3273; day 14 males: p=0.3094, r=0.4128) or SOD2 (day 7 females p=0.271, r=0.4435; day 7 males: p=0.1865, r=0.52; day 14 females p=0.4813, r=0.322; day 14 males: p=0.2546, r=0.4573).

Running increased SOD1 gene expression in DRG from injured and sham male and female rats at day 7 (sham and CCI male and females p<0.0001), but not day 14 post surgery (p=0.0905). Running had no effect on SOD2 expression in DRG (day 7 p=0.7585; day 14 p=0.1688) or SOD1 or SOD2 expression in spinal cord (SOD1 day 7: p=0.4640, day 14: p=0.0439; SOD2 day 7: p=0.9674, day 14: p=0.1441) (data not shown).

HO-1 was upregulated by CCI relative to sham operated rats in the sciatic nerve at day 7 and at day 14 in all groups (p<0.0001, Fig. 5 E,F). Running did not alter HO-1 expression in the sciatic nerve. At day 7, prior running increased expression of HO-1 in DRG of sham operated males, compared to sedentary controls (p=0.0201) (data not shown). Contrary to HO-1, Heme Oxygenase 2 (HO-2) was decreased by CCI surgery in the sciatic nerve at day 7 and 14 relative to sham operated rats in all groups (p<0.0001, Fig.5 G,H). Running did not alter HO-2 expression in the sciatic nerve. HO-2 was not altered by surgery or running in the DRG (day 7: p=0.6054, day 14: p=0.9999) (data not shown). These results show that running increases antioxidant capacity in sciatic nerve and DRG, pointing to increased scavenging of superoxide as a probable mechanism for reduction in peroxynitrite.

### Nrf2 activation during running preconditioning is required for prevention of neuropathic pain

Nrf2 is considered the master regulator of the antioxidant response, and is responsible for transcription of superoxide dismutases and heme oxygenases (Cuadrado et al., 2019; Schmidlin et al., 2019; Grace et al., 2021). We therefore investigated whether Nrf2 nuclear translocation, a marker of activation, was enhanced by prior running.

Running increased sciatic Nrf2 nuclear translocation at day 7 post surgery in both sexes in sham operated rats (male and female sham: p=0.0038) (Fig. 6A), and at day 14 post surgery in both sexes of CCI rats (male and female CCI p=0.0003) (Fig. 6B). CCI surgery increased expression of Nrf2 relative to sham in rats with prior running at day 14 post CCI (male and female runners: p<0.0001), but there was no effect of CCI surgery on Nrf2 expression at day 7. There was no effect of CCI or running on Nrf2 expression in DRG (data not shown). There were no correlations between wheel running distance or speeds and expression levels of Nrf2 (day 7 females: p=0.9912, r=0.0035; day 7 males: p=0.1007, r=-0.5205; day 14 females: p=0.4385, r=0.2608; day 14 males: p=0.3171, r=-0.3329).

**Figure 6.**
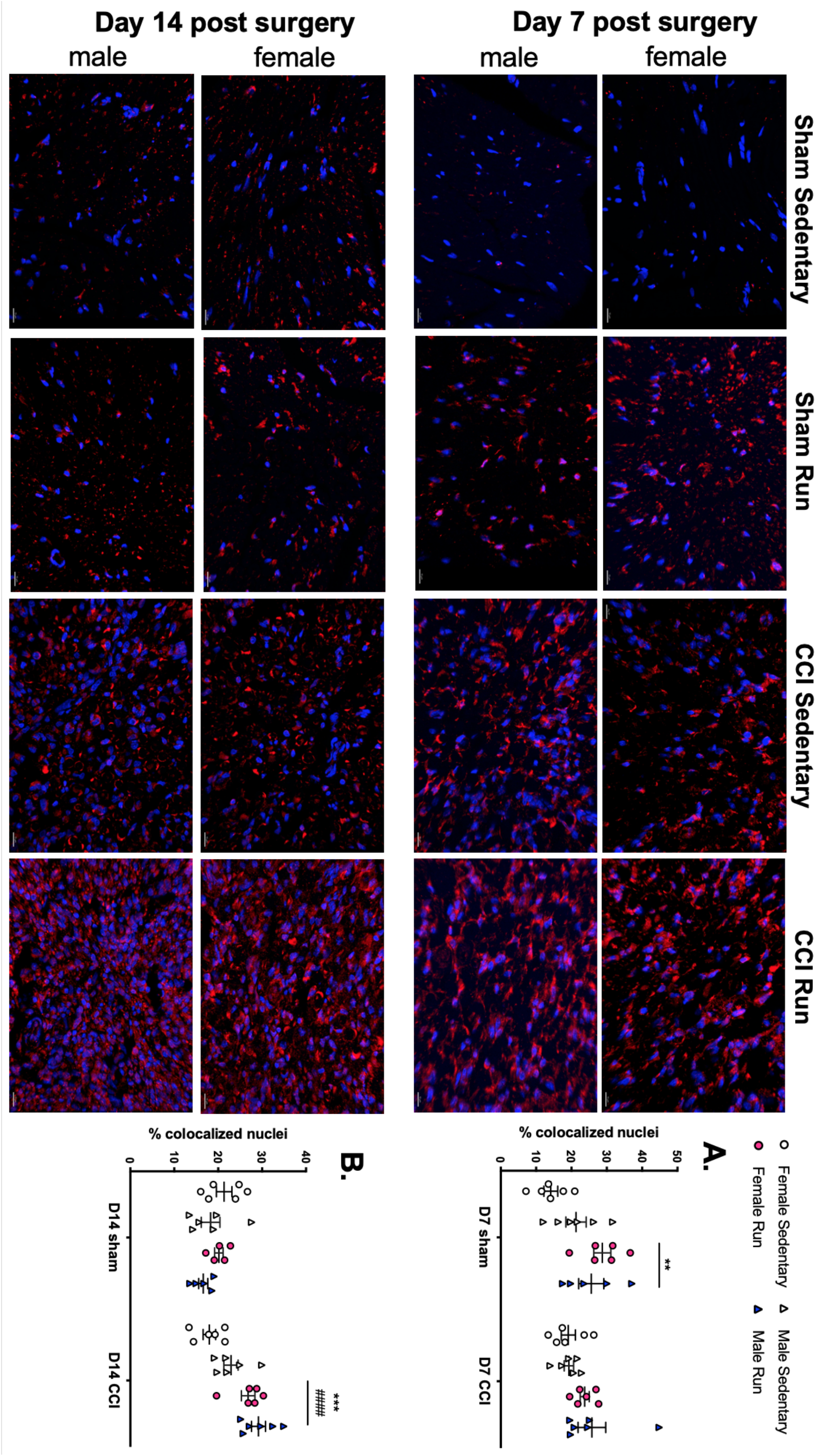
Nrf2 nuclear translocation was quantified in the sciatic injury site at day 7 **(A)** and 14 **(B)** post surgery. Scale bar = 15um. **p<0.01, ***p<.001, CCI + sedentary vs. CCI + run. ####p<.0001, sham vs. CCI.

Given that 3 weeks of prior voluntary wheel running did not protect against subsequent neuropathic pain (Fig. 2), we tested whether this short duration of prior running was also sufficient to enhance Nrf2 activation. In contrast to 6 weeks of prior voluntary wheel running, 3 weeks of prior running did not increase nuclear translocation of Nrf2 in sciatic nerve at day 14 (Fig. 7; p=0.9181). Nrf2 activation in the sciatic was therefore correlated with prevention of neuropathic pain.

**Figure 7.**
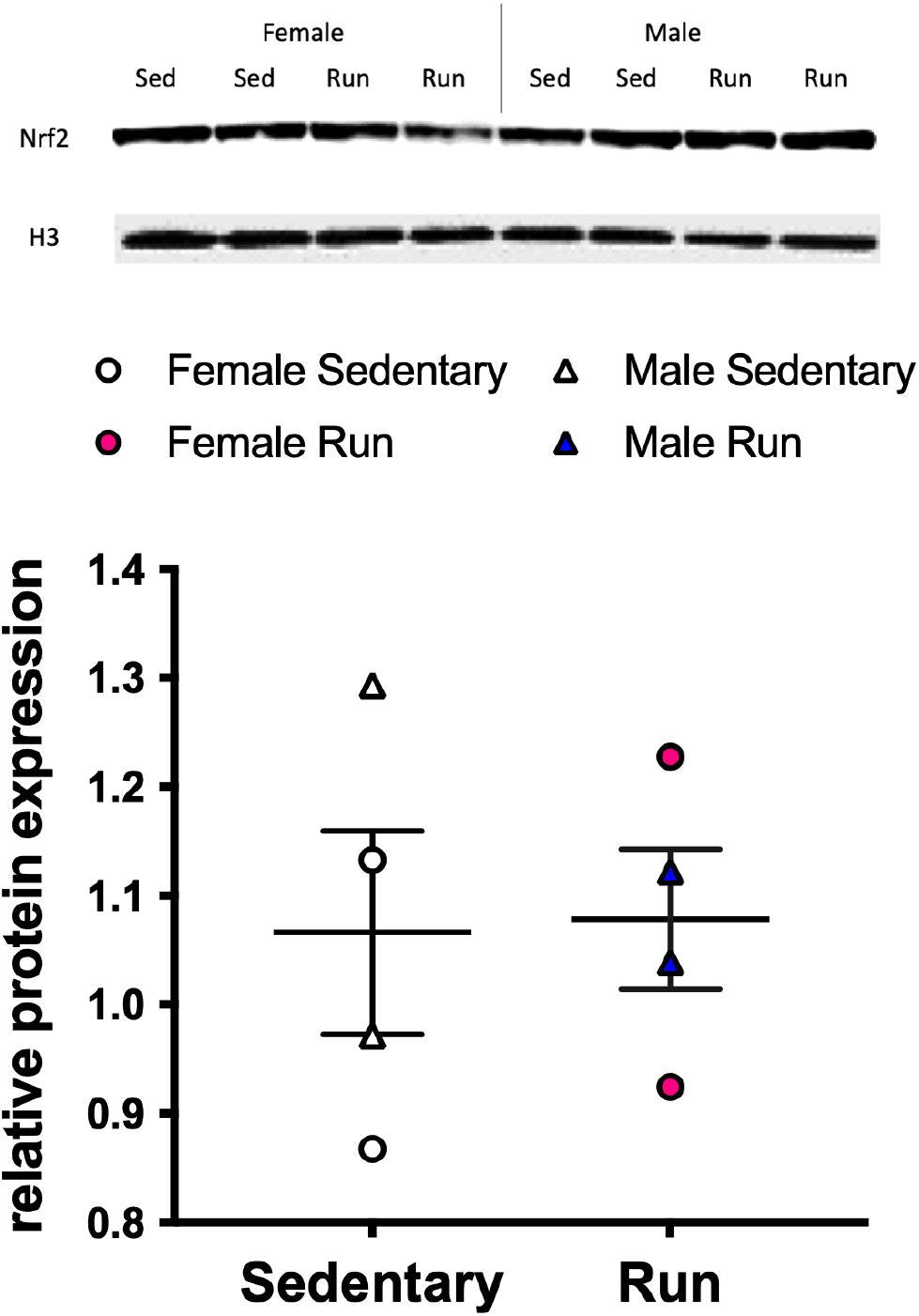
Nrf2 nuclear fractions isolated from sciatic nerve injury site at day 14 post surgery were quantified in rats that had run for 3 weeks prior to CCI vs sedentary for 3 weeks prior to CCI. Sed = sedentary. H3 = histone 3 loading control.

Our final experiment tested whether activation of Nrf2 during prior running was necessary for prevention of neuropathic pain. Nrf2 was inhibited for the 6 weeks of wheel running by administering trigonelline in the drinking water. Trigonelline did not alter running activity, compared to regular drinking water (data not shown). The protective effect of prior voluntary wheel running was abolished in rats that were treated with trigonelline, compared to vehicle (Fig. 8; treatment x time: F_9,60_ =10.74, p<0.0001; treatment F_3,20_= 149.5, p<0.0001; time F_2.67,53.44_= 15.96, p<0.0001), from weeks 2-4 after injury (p<0.05). These data show that activation of the antioxidant master-regulator Nrf2 during the 6 weeks of voluntary wheel running is necessary to confer protection from neuropathic pain after peripheral nerve injury.

**Figure 8.**
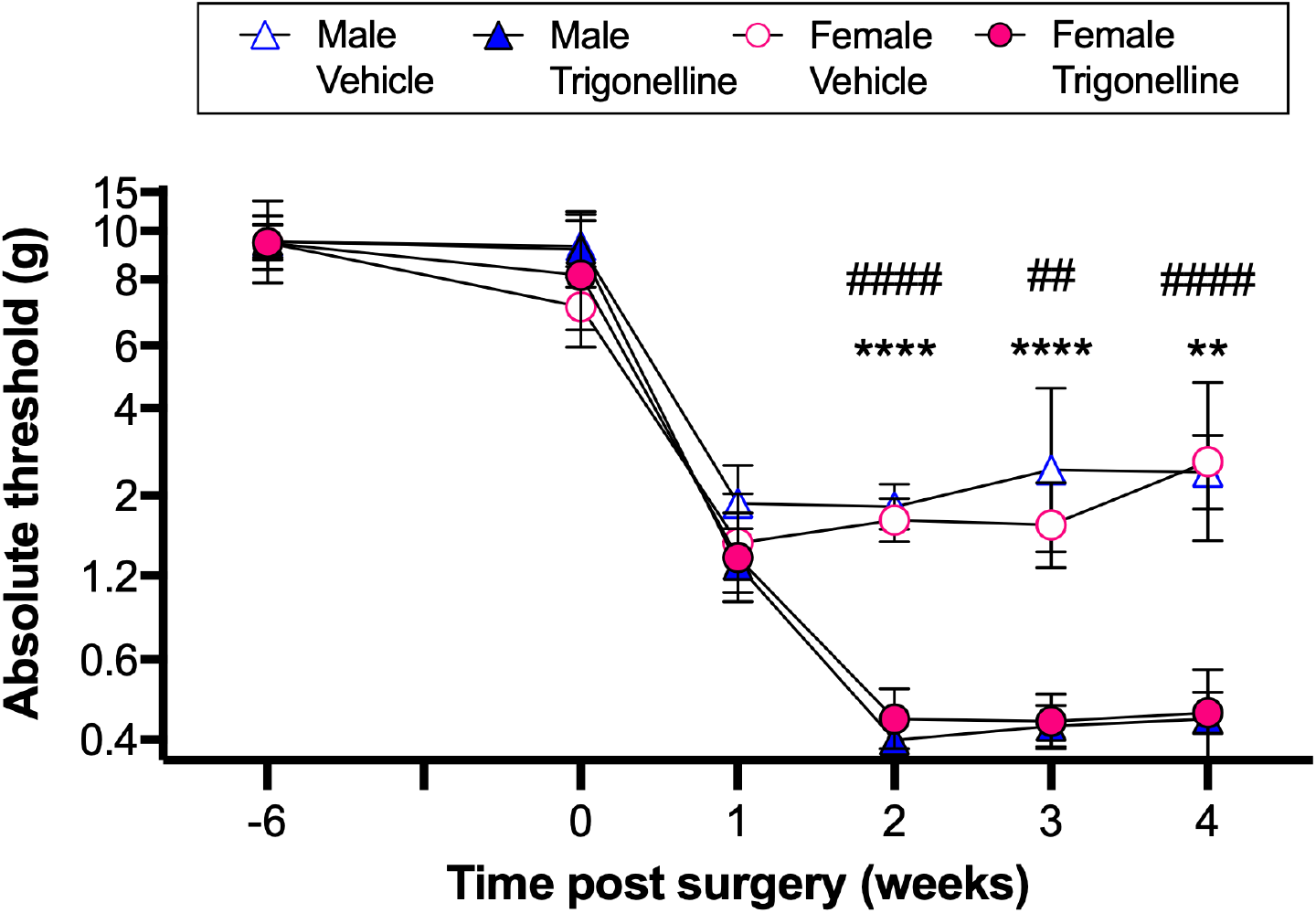
Trigonelline, an inhibitor of Nrf2, was given during the 6 weeks of voluntary wheel running prior to CCI surgery in male and female rats. Rats were tested for mechanical allodynia for four weeks following surgery. Running behavior was not altered by trigonelline administration (data not shown). **p<0.01, ****p<.0001, female trigonelline vs female vehicle, ##p<0.01, ####p<.0001, male trigonelline vs male vehicle.

## Discussion

We report that neuropathic pain induced by peripheral nerve injury is prevented by 6 weeks of prior voluntary wheel running across both sexes of adult rats. Attenuated neuropathic pain was associated with reductions in peroxynitrite—a pronociceptive reactive product of nitric oxide and superoxide (Salvemini et al., 2011)—at the site of peripheral nerve injury. Our data suggest that prior wheel running prevents formation of peroxynitrite by increasing scavenging of the superoxide; SOD1 and SOD2 expression was increased, whereas expression of the enzymatic generators of superoxide and nitric oxide, or the nitric oxide scavengers (HO-1, HO-2) were not influenced by wheel running. Prevention of neuropathic pain was further associated with activation of the antioxidant master regulator, Nrf2. Six weeks of prior voluntary wheel running increased Nrf2 nuclear translocation at the sciatic nerve injury site; in contrast, 3 weeks of prior wheel running, which failed to prevent neuropathic pain, had no effect on Nrf2 nuclear translocation. Finally, we demonstrated that the preventative effects of voluntary wheel running are mediated by Nrf2, as neuropathic pain was fully expressed across both sexes when Nrf2 activation was blocked during the running phase.

We have discovered that exercise prior to peripheral nerve injury protects against subsequent neuropathic pain by activating Nrf2. Repeated bouts of exercise acutely increase ROS/RNS production by skeletal muscle and leukocytes, among other cells (Margaritelis et al., 2020). As an adaptive response, these species activate Nrf2 across both sexes, resulting in production of antioxidants (Done and Traustadottir, 2016; Grace et al., 2021). Such effects are apparent in our data, as prior voluntary wheel running also altered expression of antioxidants in sham-operated animals. We propose that such preconditioning by prior exercise facilitates a rapid and robust antioxidant response to traumatic peripheral nerve injury, restraining secondary damage and other pronociceptive effects of ROS/RNS. In support, we have previously shown that prior voluntary wheel running reduces DRG levels of ATF3, a marker of neuronal injury (Grace et al., 2016b). This proposal is also consistent with the sexually monomorphic anti-neuropathic function of Nrf2 (Rosa et al., 2008; Wang and Wang, 2017; Ferreira-Chamorro et al., 2018; Yang et al., 2018; Li et al., 2020; Grace et al., 2021). It is unknown whether Nrf2 is similarly engaged in reversal of established neuropathic pain by exercise (Cobianchi et al., 2010; Stagg et al., 2011; Li and Hondzinski, 2012; Groover et al., 2013; Benson et al., 2015; Bobinski et al., 2015; Grace et al., 2016b; Chhaya et al., 2019).

Although a role for Nrf2 in the antinociceptive effects of exercise has not been previously explored in pain states, other groups have evaluated regulation of antioxidants by exercise. In models of neuropathic pain induced by traumatic nerve injury or experimental autoimmune encephalomyelitis, reversal paradigms of exercise increased the reduced glutathione to oxidized glutathione (GSH/GSSG) ratio in the lumbar spinal cord, and antioxidant activity and capacity in serum (Benson et al., 2015; Mifflin et al., 2017; Safakhah et al., 2017; Rostami et al., 2020). In our study, voluntary wheel running increased expression of SOD1 and SOD2 at the sciatic nerve injury site and DRG, but not lumbar spinal cord. While HO-1 was upregulated by CCI, there was no interaction with voluntary wheel running at the timepoints assessed. However, evaluation of acute timepoints after nerve injury is necessary to rule out any interactions. We did not detect CCI-induced increases in any antioxidant genes in the lumbar spinal cord, but the literature is mixed on this subject as antioxidant responses are temporally- and injury-specific (Grace et al., 2016a).

The reduction in peroxynitrite by prior voluntary wheel running is consistent with its well-established role in neuropathic pain. Pro-nociceptive mechanisms include mitochondrial dysfunction in sensory neurons via SOD2 nitration and lipid peroxidation, and induction of proinflammatory cytokines (Matata and Galinanes, 2002; Salvemini et al., 2011; Doyle et al., 2012; Little et al., 2012; Janes et al., 2013; Grace et al., 2014b; Grace et al., 2016a; Grace et al., 2021). It is therefore possible that the attenuated pro-inflammatory signaling at the site of nerve injury, observed previously with 6 weeks of prior voluntary wheel running (Grace et al., 2016b), is a downstream consequence of reduced peroxynitrite levels. CCI *per se* induced a strong increase peroxynitrite at the site of nerve injury, aligning with other studies of traumatic nerve injury (Liu et al., 2000). However, we only detected modest increases in DRG, and no change spinal cord, in contrast to models of diabetic and chemotherapy-induced neuropathy (Obrosova et al., 2007; Vareniuk et al., 2007; Doyle et al., 2012; Little et al., 2012; Janes et al., 2013). Thus, the production of peroxynitrite along the pain neuraxis may be dependent on the nature of the injury to the sciatic nerve. It would also be instructive to determine the influence of exercise on additional ROS/RNS that are elevated at other sites along the pain neuraxis after peripheral nerve injury.

This study also deepens our understanding of the parameters of prior voluntary wheel running that confer protection from neuropathic pain; six weeks is sufficient, but three is not. It is possible that the decay of exercise-induced Nrf2 activation after injury is slowed as the duration of exercise increases. Future studies could therefore test whether the magnitude of allodynia prevented by prior running could be further increased by extending the exercise period, or if there is a threshold. As in other conditions (Greenwood et al., 2005a; Greenwood et al., 2005b), these results further rule out acute effects of running (e.g., hormonal) as an explanation for suppression of neuropathic pain. Our data further revealed that weekly distances traveled or maximum running speeds had no correlation with attenuation of allodynia or other biochemical measures, despite large inter-individual variability and sex differences in these running parameters. These data suggest that there may be a low threshold in the weekly distances required for protection from neuropathic pain. This is supported by previous work showing that aged animals running as little as 700 m a week were protected from infection-induced memory impairments (Barrientos et al., 2011).

In sum, our present study offers a mechanism by which prior voluntary wheel running prevents subsequent neuropathic pain across both sexes. Six weeks of prior exercise increases expression of SOD1 and SOD2 via activation of Nrf2 in the sciatic nerve injury site, attenuating formation of peroxynitrite that otherwise drives nociceptive hypersensitivity. This study provides further evidence that physical activity may be an effective strategy to prevent severe neuropathic pain (Landmark et al., 2011; Landmark et al., 2013).

## Acknowledgements

This work is supported by National Institutes of Health grants R01 AT009366 (L.R.W.), RF1 NS113840 (P.M.G.), and P30 CA016672 (MD Anderson Cancer Center).

